# Predicting subject traits from brain spectral signatures: an application to brain ageing

**DOI:** 10.1101/2023.11.02.565261

**Authors:** Cecilia Jarne, Ben Griffin, Diego Vidaurre

**Affiliations:** Departamento de Ciencia y Tecnologia de la Universidad Nacional de Quilmes, Bernal, Buenos Aires, Argentina; CONICET, Buenos Aires, Argentina; Center of Functionally Integrative Neuroscience, Department of Clinical Medicine, Aarhus University, Aarhus, Denmark; Nuffield Department of Clinical Neurosciences, Oxford University, Oxford, UK; Oxford Centre for Human Brain Activity, Department of Psychiatry, Oxford University, Oxford, UK

**Keywords:** Kernel Mean Embedding Regression, Brain age, EEG, Machine Learning, Kernel methods, Maximum Mean Discrepancy

## Abstract

The prediction of subject traits using brain data is an important goal in neuroscience, with relevant applications in clinical research, as well as in the study of differential psychology and cognition. While previous prediction work has predominantly been done on neuroimaging data, our focus is on electroencephalography (EEG), a relatively inexpensive, widely available and non-invasive data modality. However, EEG data is complex and needs some form of feature extraction for subsequent prediction. This process is sometimes done manually, risking biases and suboptimal decisions. Here we investigate the use of data-driven kernel methods for prediction from single-channels using the the EEG spectrogram, which reflects macro-scale neural oscillations in the brain. Specifically, we introduce the idea of reinterpreting the the spectrogram of each channel as a probability distribution, so that we can leverage advanced machine learning techniques that can handle probability distributions with mathematical rigour and without the need for manual feature extraction. We explore how the resulting technique, Kernel Mean Embedding Regression, compares to a standard application of Kernel Ridge Regression as well as to a non-kernelised approach. Overall, we found that the kernel methods exhibit improved performance thanks to their capacity to handle nonlinearities in the relation between the EEG spectrogram and the trait of interest. We leveraged this method to predict biological age in a multinational EEG data set, HarMNqEEG, showing the method’s capacity to generalise across experiments and acquisition setups.

## 1. Introduction

Predicting behavioural and cognitive traits from brain data in a way that generalises to unseen subjects and is robust to acquisition idiosyncrasies is important because it can offer objective measures to otherwise elusive neurobiological constructs (Haynes and Rees (2006)). A widely studied example is brain age. While measuring actual age is a straightforward task, the concept of brain age provides a marker of mental health by quantifying how much a subject’s brain appears to have aged with respect to the population average; that is, a predicted age that is lower than the individual’s chronological age, for example, may indicate that the person has a brain that appears younger than expected for their actual age (Franke and Gaser (2019); Smith et al. (2019)).

Much of the work on the prediction of subject traits (such as age) from brain data has been done with resting-state fMRI data (Dosenbach et al. (2010)). Since we cannot straightforwardly predict from the raw data, an intermediate representation is typically used for prediction. In the case of fMRI, this is often a simple description of functional connectivity (Rosenberg et al. (2016)) or some model of brain dynamics (Liegeois et al. (2019); Vidaurre et al. (2021); Ahrends et al. (2023)). Here, we focus on EEG, a considerably less costly technique. As an intermediate representation, we consider the EEG frequency spectrum, reflecting neural oscillations that are well-known correlates of different behavioural and cognitive states (Buzsáki and Draguhn (2004)). How to meaningfully extract features for prediction from an EEG spectrum is an open question. Some current efforts are based on manual feature extraction (Al Zoubi et al. (2018); Engemann et al. (2022)); or are based on models that need to be estimated from raw data and whose properties depend on the choice of configuration and hyperparameters (Vidaurre et al. (2013)).

In this paper, we present a method to predict from single-channel spectrograms, i.e. with no need of the raw data. In contrast to other works that make the predictions on whole brain’s signals (Dimitriadis and Salis (2017); Vandenbosch et al. (2019); Sabbagh et al. (2019)), sometimes with complex methods such as deep learning (Khayretdinova et al. (2022)), our approach has not only benefits in terms of computational simplicity but also for interpretation, as it allows us to compare the predictive power across sensors or brain regions. Specifically, with no prior predefinition of frequency bands or any manual feature engineering besides basic preprocessing, we propose the idea of interpreting the EEG spectrogram as a probability distribution. This way, we can fully leverage all the powerful machinery of kernel learning on probability distributions. We consider Kernel Mean Embeddings (KME), a technique used to construct a representation of the data in a high-dimensional feature space (Smola et al. (2007); Iyer et al. (2014); Borgwardt et al. (2006)). The KME technique maps joint, marginal and conditional probability distributions to vectors in a high (or even infinite) dimensional feature space that completely characterises the distribution (Fukumizu et al. (2011)). Building upon KME, we used the Maximum Mean Discrepancy (MMD) (Smola et al. (2007)), a distance metric defined on the space of probability measures (here, EEG spectrograms) in combination with kernel ridge regression (Saunders et al. (1998)). We refer to our approach as Kernel Mean Embedding Regression (KMER), which we compare to Kernel Ridge Regression (KRR) and Ridge Regression (RR).

We demonstrate the superior performance of the kernel approaches (both KMER and KRR) for age prediction. Using a multisite, public resting-state EEG dataset with a very wide distribution of age (HarMNqEEG, Li et al. (2022)), we show that these lead to better predictions, which can be projected on the EEG scalp for interpretation. Focusing on KMER, we show that parietal sensors are the most accurate in predicting age, with slightly greater accuracy in men. We also showed that the predictions generalise well across experiments and acquisition sites, even considering the large differences in age distribution across sites (a well-known problem in machine learning referred to as prior shift). Overall, by demonstrating its predictive capacity and interpretability, kernel methods can help unveil insights about brain age or other neurobiological constructs.

## 2. Results

### 2.1. Kernel mean embedding regression

We developed KMER, a novel method to predict subject traits from EEG spectrograms; see Methods and Figure 1. In short, KMER is based on the idea of interpreting EEG spectrograms as probability distributions so that techniques based on kernel learning of distributions are readily applicable. By forming a subject-by-subject kernel matrix of similarities between subjects (using a Gaussian kernel; see Methods), this technique allowed us to derive predictions for each EEG channel that can naturally accommodate non-linearities in a data-driven way without the need for manual feature extraction.

**Figure 1:**
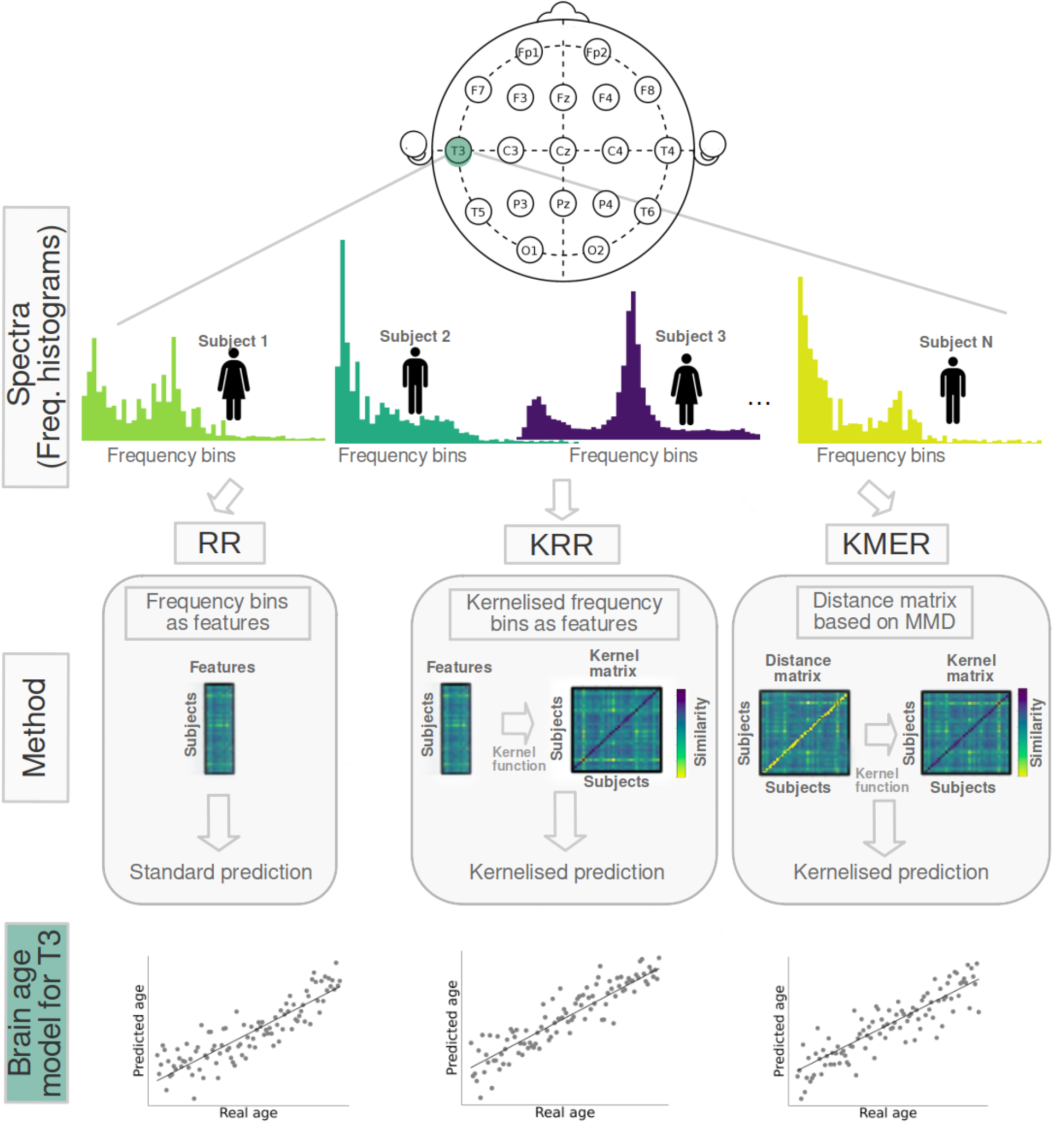
General workflow of the analysis. We used the HarMNqEEG dataset collected across multiple sites (batches), which contains rescaled power densities per sensor and participant, as well as information on gender and age. We considered three prediction approaches, which we run separately per EEG sensor: (i) Ridge Regression (RR) using the power estimates at each frequency bin of the spectrogram as features; (ii) Kernel Ridge Regression (KRR), a kernelised version of RR (based on a non-linear radial basis function kernel); and (iii) our Kernel Mean Embedding Regression (KMER) approach, where we interpreted the power spectral estimates across bins as probability distributions so that we can leverage the mathematical machinery of kernel learning on probability measures for prediction.

### 2.2. Comparison of performance between methods

First, compared KMER to: (i) RR, a regularised, linear regression method that uses the frequency bins’ power as features; and (ii) KRR, the kernelised version of RR, which allows the modelling of nonlinearities by the use of an appropriate kernel. Our experiments were based on the prediction of age in the HarMNqEEG dataset (Li et al. (2022)), a multibatch, multicountry resting-state EEG dataset of 1966 subjects encompassing people across the entire lifespan (excluding the very youngest and using only subjects between 5 and 97 years old); see Figure S1 and Table S1 (SI) for some basic statistics about the dataset in terms of age, gender, and geography. The performance of the methods was assessed using cross-validation, such that the acquisition batches were never split across folds. The hyperparameters (described in Methods) were chosen using nested cross-validation for all approaches. We assessed the performance of the three methods per EEG channel using two measures of accuracy: prediction explained variance (*R*^2^) and mean absolute error (MAE), which offer complementary information (Engemann et al. (2022)).

Figure 2 summarises the results per EEG channel. Figure 2A illustrates an example of age vs. predicted age for a given channel (T3). Figure 2B and 2C show a comparison of the three methods across channels in terms of MAE and explained variance, respectively. As shown, MAE oscillates between 11 and 12.5 years approximately for all channels. There is a larger variation in terms of explained variance, ranging from *R*^2^ = 0.24 for sensor Fp1 to *R*^2^ = 0.44 for sensor C3. KRR and KMER outperform RR for all channels, and KMER outperforms KRR only for some channels. The advantage of KRR and KMER over RR is highly significant (p-value = 1 *×*10 − 5; permutation testing), but the advantage of KMER over KRR is not significant (p-value=0.093). This improvement of the kernel methods over RR is likely due to the presence of nonlinearities in the relation between EEG spectrograms and age. Results for the linear and polynomial kernel (referring to the kernel used to construct the kernel matrix, *f* (·); see Methods) are shown in Figures S2 and S3 (SI); these performed worse than the radial basis function (Gaussian) kernel.

**Figure 2:**
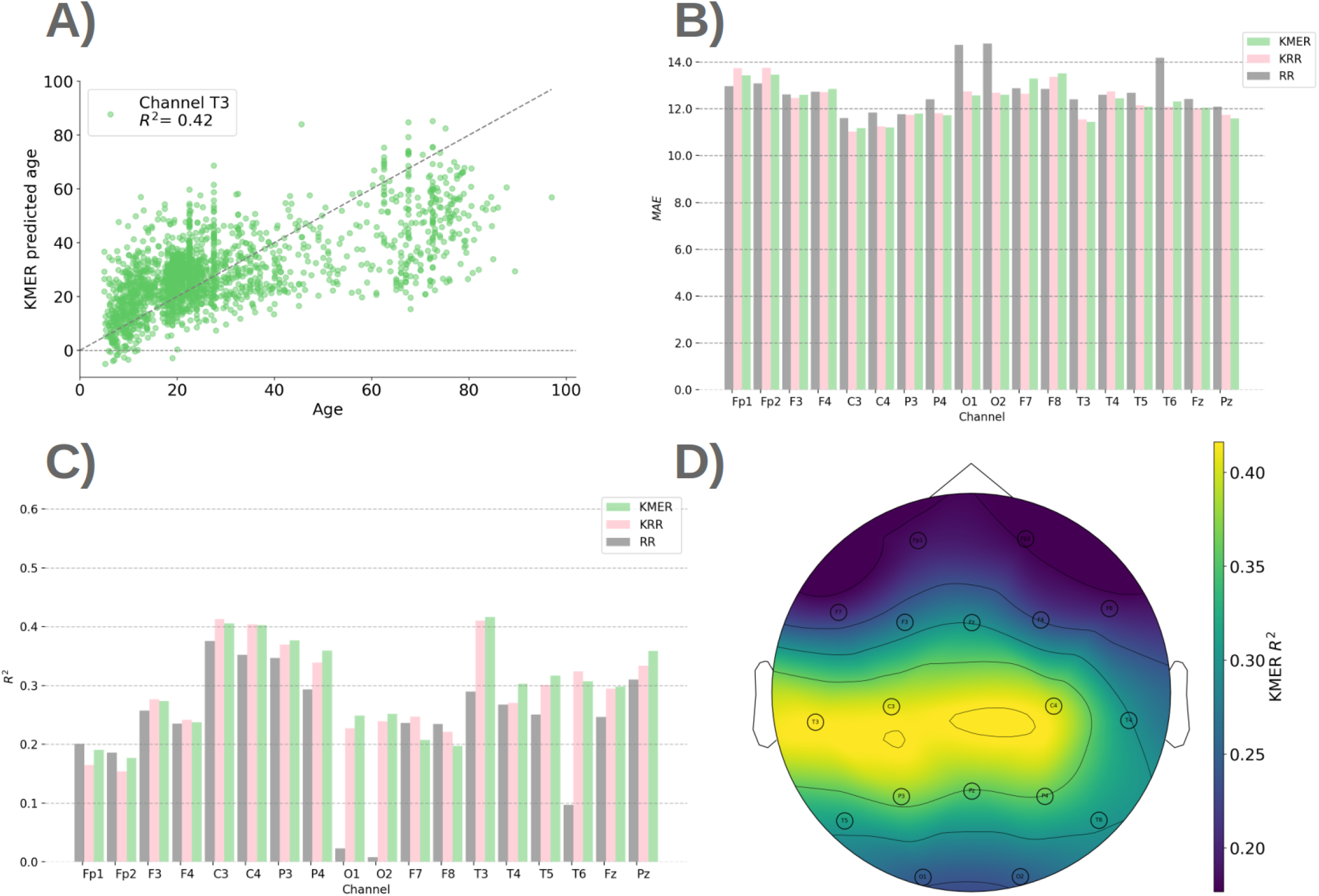
Comparison between the methods. **A)** Illustration of predicted vs. real age for example channel T3 for KMER. **B)** Mean Absolute Error (MAE) per channel for KMER, RR and KRR. **C)** Explained variance *R*^2^. **D)** *R*^2^ projected on sensor space for KMER.

An important observation, as seen in Figure 2A, is that age differences are less well predicted at older ranges. To try to address this problem, we considered the prediction of age in log space (such the differences in the younger are amplified). As observed in Figure S4, we obtained moderate improvements in *R*^2^ and MAE.

A posthoc bias correction, where we removed the age dependence on the residual in a second step using linear regression (Smith et al. (2019)), however resulted in a more substantial improvement; see Figure S5. This correction is performed by training a linear regression model on the residuals of the training data to decrease the bias of the predictions in the testing data, typically improving performance across metrics like *R*^2^ and MAE. Specifically, we first calculate the residuals of the training data by subtracting the predictions from the actual training values. Then, we fit a linear regression model, where the chronological age is the independent variable and the residual is the dependent variable. This makes the (new) residual and age orthogonal (Smith et al. (2019)). Here, the correction is incorporated to the original KRR/KMER predictions to generate bias-corrected predictions.

We also studied the effect of normalising each frequency bins across subjects, such that the variance is equal for all bins. As shown in Figure S7, the results did not change substantially.

Finally, Figure 2D shows the explained variance in sensor space for KMER (MAE has a similar but inverted topography; not shown). We can observe that the parietal sensors are the most predictive of chronological age, whereas prefrontal and occipital sensors are the least predictive.

### 2.3. Sex differences

Females and males have previously been shown to exhibit differences in their ageing trajectories (Hägg and Jylhävä (2021)). Here, using KMER, we investigated the differences between sexes in prediction accuracy across channels for 884 females and 905 males (for 137 individuals, sex is non-specified).

Figure 3 presents the results. Figure 3A and 3B show example scatter plots between age and predicted age per sex. Figure 3B and 3C show *R*^2^ across sensors in both barplot and sensor space format. Analogously, Figure 3D and 3E show MAE per sex. As observed, males generally exhibited a higher prediction accuracy than females (p-value=0.002), potentially suggesting that their pace of biological change corresponds more closely to chronological age; see Figure S6 for a side-by-side comparison.

**Figure 3:**
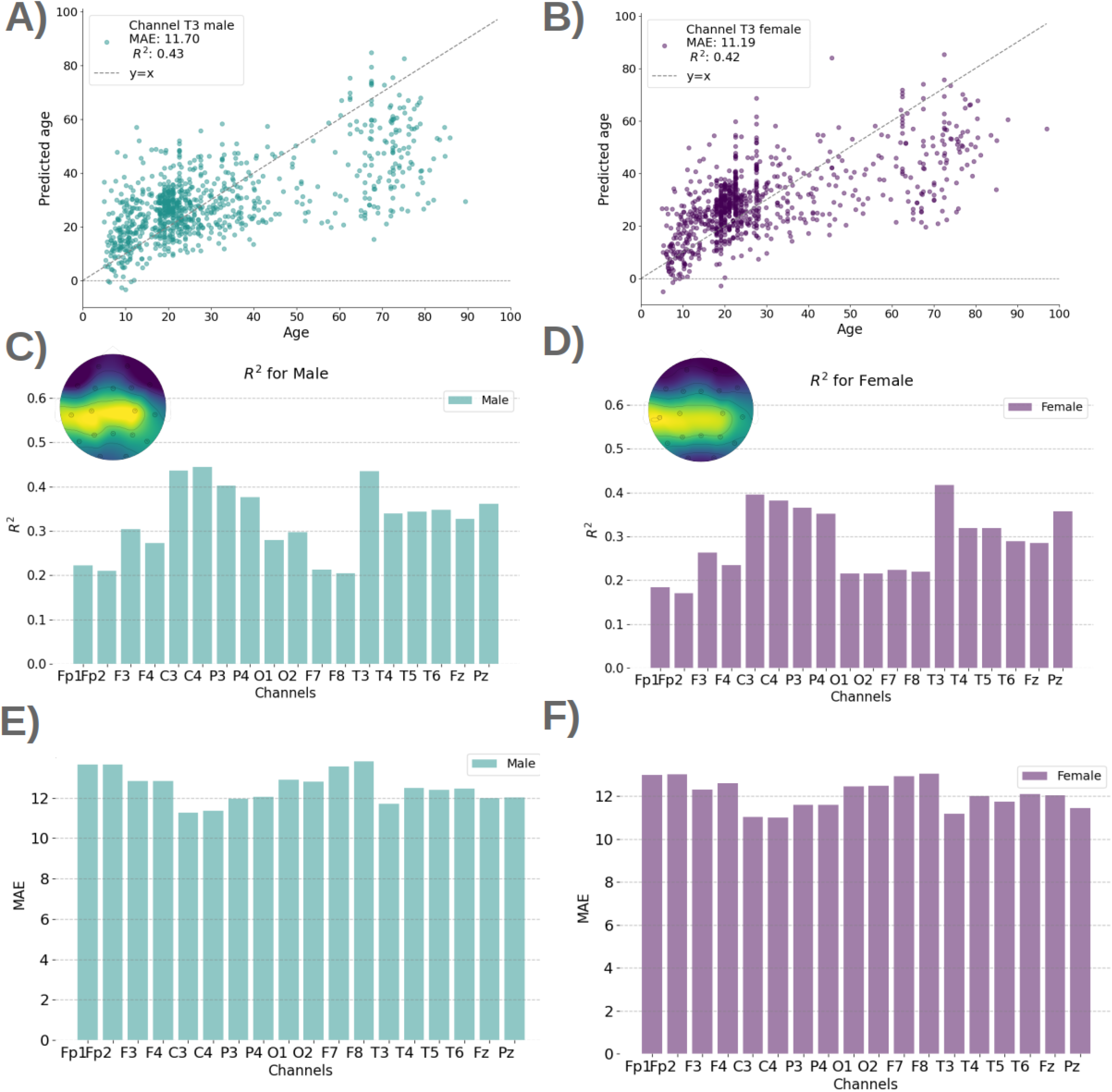
Sex differences (left panels, male; right panels, female). **A)** and **B)** Predicted vs. real age per sex for example channel T3; results are consistent for all channels (not shown). **C)** and **D)** Explained variance *R*^2^ per sex and channel, in barplot format and in a topographic map. **E)** and **F)** MAE per sex and channel.

### 2.4. Performance variations across scanning sites

The HarMNqEEG dataset encompasses a wide range of age groups across various sites or geographical locations (14 different sites/experiments), each with a different number of individuals and specific age distributions; see Figure S1 and Table S1. Despite efforts made by the data curators to ensure homogeneity, it is possible that differences in practice and instrumentation between the sites leak into the data. In our reported results so far, sites were never split across cross-validation folds; that is, when predicting the age of subjects in a given site (e.g. Colombia), none of the subjects of that site were part of the training data. Given that substantial differences in age distribution between sites may potentially coexist with other between-site differences (e.g. instrumentation), this approach was performed to ensure that the reported prediction accuracies were purely reflective of age and not mixed with other factors. However, this conservative approach makes the prediction more difficult, inducing what is known in the machine learning literature as a prior shift (Kouw (2018)), where the distribution of the dependent variable (here, age) changes between training and test.

To illustrate the issue, Figure 4A shows predicted age vs. chronological age with colours indicating stratification by site, with two different perspectives: 3D to better appreciate each site individually and 2D to compare the sites side-by-side. Here, we can observe substantial differences between sites in both accuracy and age distribution. Figure 4B shows distributions of delta across sites, defined as the difference between predicted and chronological age (Franke and Gaser (2019); Smith et al. (2019)). When compared with Figure S1, where we show the age distributions explicitly, we can observe that the distribution of errors is contingent on the age range within each group, underestimating the age of older individuals (see Switzerland as an example, which has the oldest population in the data set). The observed pattern is consistent across different channels (not shown).

**Figure 4:**
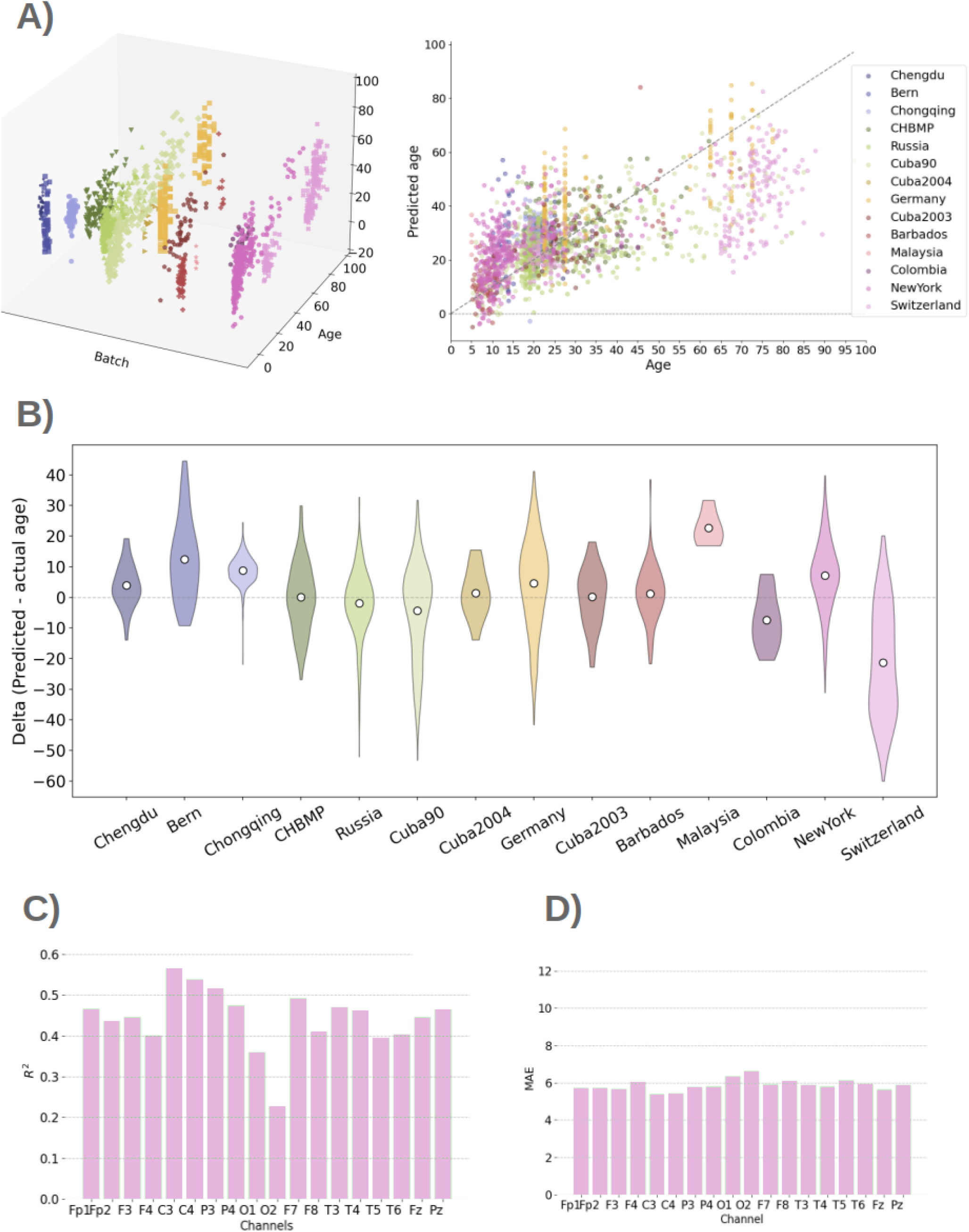
Differences between sites and their effect on the prediction. **A)** Predicted vs. real age in 3D (left) and 2D (right), separated by site (batch). **B)** Distribution of delta (defined as predicted minus real age) per site. **C** and **D)** *R*^2^ and MAE where training and testing come from a single site (New York).

We compared the cross-site to a within-site prediction, where we ran a cross-validated prediction within one given site without using data from the other sites. While this can only be done effectively for sites with a sufficient number of subjects, we observed substantial increases in accuracy in this type of prediction. Figure 4CD shows the case of New York, where, the explained variance went up to 0.63 and MAE became as small as 5.1.

Overall, these results demonstrate the challenges of predicting across sites when the age distributions is very different, but also that KMER and KRR can still produce reasonable accuracies even in this adverse situation.

### 2.5. Comparison with other studies

To further compare with existing work on age prediction, Table 1 presents the prediction accuracies from other studies that performed age estimation from EEG data, using other methodological approaches and data sets. Considering that our method operates at the single channel level, that it only has access to the spectrograms, and that our reported accuracies are primarily across sites (which, as mentioned, is harder), the present accuracies fare well in relation to those reported elsewhere.

**Table 1:**
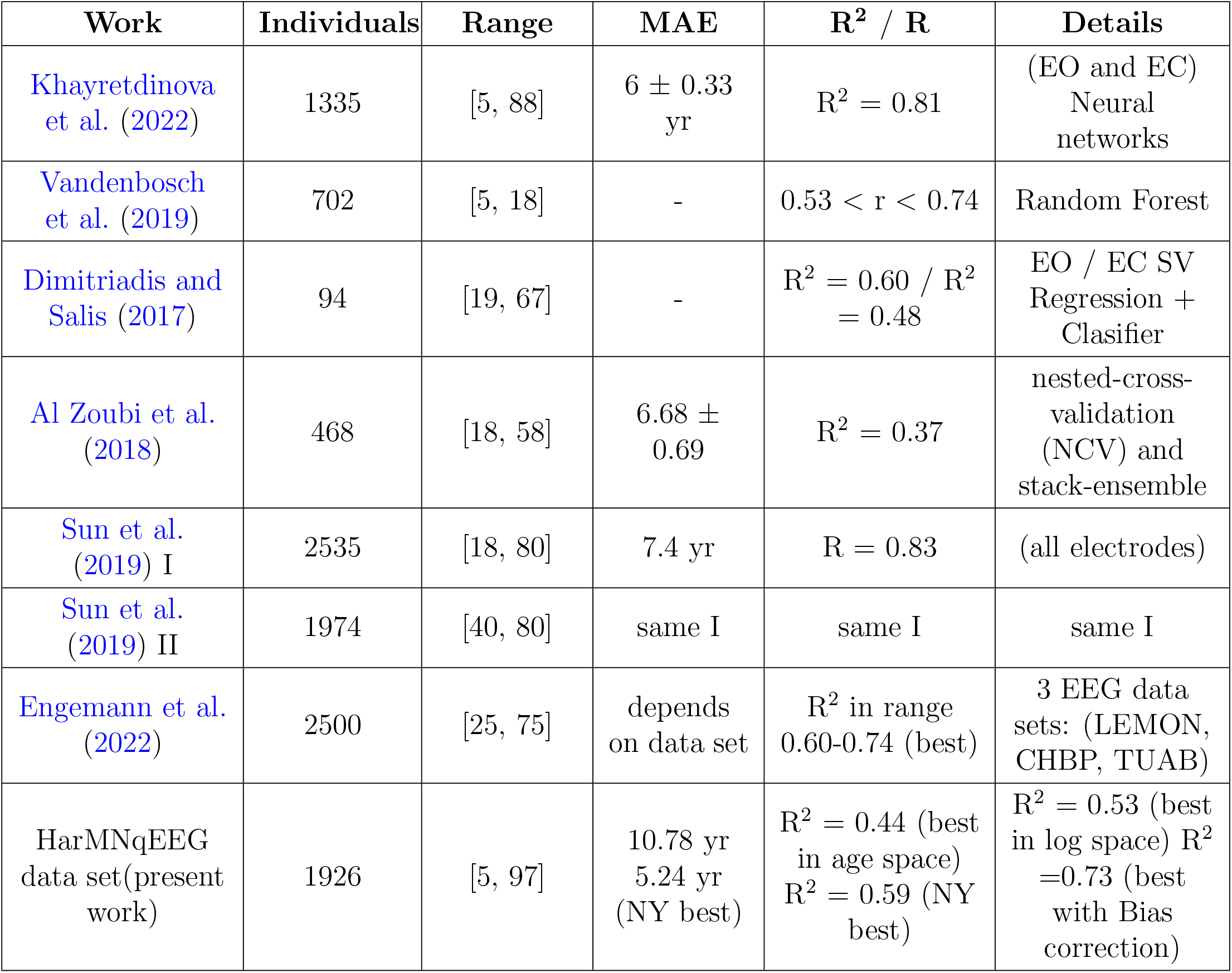
Other results from the literature of age estimation using EEG. These figures were obtained using different methods, age ranges and datasets. EO and EC, respectively, denote eyes open and eyes closed. The results are compared with ours for the best channel in age space, for NY, in log space and using bias correction (See Figure S5).

| Work | Individuals | Range | MAE | $R^2$ / R | Details |
| --- | --- | --- | --- | --- | --- |
| Khayretdinova et al. (2022) | 1335 | [5, 88] | $6 \pm 0.33$ yr | $R^2 = 0.81$ | (EO and EC) Neural networks |
| Vandenbosch et al. (2019) | 702 | [5, 18] | - | $0.53 < r < 0.74$ | Random Forest |
| Dimitriadis and Salis (2017) | 94 | [19, 67] | - | $R^2 = 0.60$ / $R^2 = 0.48$ | EO / EC SV Regression + Classifier |
| Al Zoubi et al. (2018) | 468 | [18, 58] | $6.68 \pm 0.69$ | $R^2 = 0.37$ | nested-cross-validation (NCV) and stack-ensemble |
| Sun et al. (2019) I | 2535 | [18, 80] | 7.4 yr | $R = 0.83$ | (all electrodes) |
| Sun et al. (2019) II | 1974 | [40, 80] | same I | same I | same I |
| Engemann et al. (2022) | 2500 | [25, 75] | depends on data set | $R^2$ in range 0.60-0.74 (best) | 3 EEG data sets: (LEMON, CHBP, TUAB) |
| HarMNqEEG data set (present work) | 1926 | [5, 97] | 10.78 yr<br>5.24 yr (NY best) | $R^2 = 0.44$ (best in age space)<br>$R^2 = 0.59$ (NY best) | $R^2 = 0.53$ (best in log space)<br>$R^2 = 0.73$ (best with Bias correction) |

## 3. Methods and Materials

The general workflow of our analysis is presented in Figure 1. In summary, we considered the spectrogram from every subject as the only input. Predictions of age were made per channel in a cross-validation fashion, where the acquisition sites (e.g. Russia) were used as folds. We used three approaches to predict age from the EEG spectrograms: RR, KRR, and KMER. For scoring prediction performance, we used explained variance (*R*^2^) and the MAE, which offer complementary information (Engemann et al. (2022)). We also computed the so-called delta, defined as the signed difference between predicted and actual age, which is often used to quantify brain age. In this Section, we describe the data set and KMER; RR and KRR are standard and their description can be found elsewhere. All code used in this paper is open and publicly available at: https://github.com/katejarne/Kernel_Max_mean_discrepancy_EEG_Age.

### 3.1 The HarMNqEEG dataset

We used EEG spectrograms from the HarMNqEEG dataset (Li et al. (2022)), which originated from a legacy dataset associated with the Cuban Human Brain Mapping initiative (Valdes-Sosa et al. (2021)). The data used in our study were collected from 9 countries, 12 EEG systems, and 14 experiments, which here we refer to as batches or sites. Overall, there are 1966 subjects, of which 40 subjects without a recorded age were excluded. Also, we discarded babies and toddlers, considering only subjects between 5 and 97 years old. For prediction, we used the rescaled raw spectrogram, such that (just like a probability distribution) the values integrate to 1.0. Additional details of the data set are described in SI, including batch names in the repository (Table S1 and Figure S1).

Recordings were taken from the 19 channels of the 10/20 International Electrodes Positioning System: Fp1, Fp2, F3, F4, C3, C4, P3, P4, O1, O2, F7, F8, T3/T7, T4/T8, T5/P7, T6/P8, Fz, Cz, Pz, where Cz was the reference electrode in some batches, while in others average-referencing was applied. We therefore discarded Cz from the analysis and performed the predictions for the other 18 channels. Data was formatted as cross-spectral matrices sampled from 1.17 to 19.14 Hz, with a 0.39 Hz resolution. The scalp EEG cross-spectrum was calculated by the data set curators using Bartlett’s method (Møller (1986)), by averaging the periodograms of more than 20 consecutive and non-overlapping segments. Because the spectra from different sites have different maximum cutoff frequencies, we used the lowest maximum cutoff frequency across sites for full compatibility. Thus, the histogram of each of the 18 channels has 49 bins corresponding to a maximum cutoff frequency of 19.13 Hz.

### 3.2. Maximum Mean Discrepancy

This section describes the mathematical foundations of KME, which the next section will use to elaborate how KME can be applied for prediction using EEG spectrograms.

Kernel mean embeddings fully characterise the distributions by mapping joint, marginal, and conditional probability distributions to vectors in a high (or infinite) dimensional feature space (Fukumizu et al. (2011); Smola et al. (2007)). This implies that any two distributions *P* and *Q* with differences in any moment are mapped to separate points on a reproducing kernel Hilbert space ℋ (Fukumizu et al. (2011)), which is essentially a space of functions. On this space, we can for instance perform classification, regression or and clustering of probability distributions, with no loss of information.

In practice, we do not have access to the distributions *P* or *Q*, so we use the available samples ***X*** = *{x*_1_, …, *x*_*n*_*}* and ***Z*** = *{z*_1_, …, *z*_*m*_*}*. The empirical estimate of the kernel mean embedding *µ*_*P*_ (i.e. the projection of the probability distribution on the reproducing kernel Hilbert space) is given by

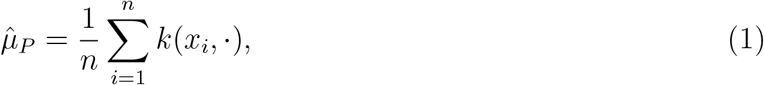

where *k*(·, ·) is a kernel function.

Now, the Maximum Mean Discrepancy (MMD), which is a distance metric established on the space of probability measures (Gretton et al. (2012)), is defined as

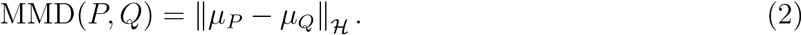

The empirical estimate of the MMD, which we use here, is given by:

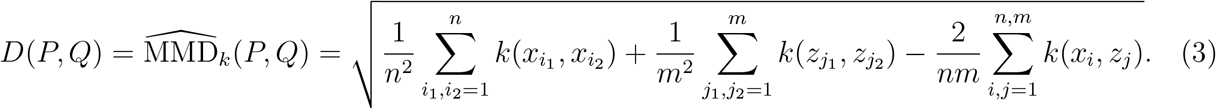

where *n* and *m* are the number of samples in distributions *P* and *Q*, respectively. In words, this formula is based in similarities between samples, and quantifies the similarities within-(first two terms) vs. between-distributions (third term). In what follows, we considered this estimate of the MMD as our distance metric between distributions (EEG spectrograms). We use a radial basis function kernel, lineal and polynomial as kernel functions.

### 3.3. Kernel Mean Embedding Regression

In this section we link the above concepts to prediction using EEG spectrograms. We interpret the EEG spectrograms (rescaled to sum up to 1.0) as probability distributions (denoted above as *P* or *Q*). We expect these to vary across age, so that we can leverage these differences for prediction. We deterministically generated samples (above, referred to as e.g. ***X***) from each rescaled spectrogram (which bin heights sum up to 1.0) so that we can compute the MMD following Equation 3. For example, for a given channel, if the bin “10Hz” has height equal to 0.2, and we assume that we have *n* = 1000 samples, then we have 0.2 *×* 1000 = 200 samples equal to 10. This way, we can apply the empirical estimator 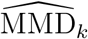 defined in Equation 3 to each pair of spectrograms in the HarMNqEEG data set, considering different kernel functions *k*(·, ·). This produced a (*N ×N*) distance matrix ***D*** per kernel, where *N* is the number of subjects. This distance matrix needs then to be converted to a kernel matrix ***K***, as required for any kernelised algorithm. For this, we use a second kernel function, denoted as *f* (·, ·), that takes a distance between two subjects to produce a quantification of similarity. Specifically, we use a radial basis function kernel,

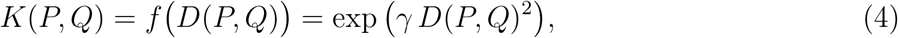

where the two subjects in question have been denoted as *P* and *Q* for continuity in the notation, and *γ* is a hyperparameter of the kernel function *f* (·). The resulting kernel matrix ***K*** can then be used to estimate prediction weights as

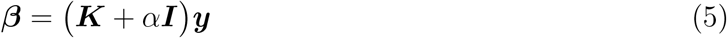

where ***y*** is the response variable (age), *α* is a regularisation hyperparameter (which we select using cross-validation) and ***I*** is the identity matrix. A prediction for an incoming subject *T* is then made as

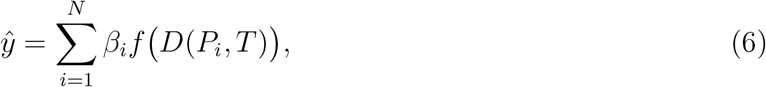

where *P*_1_, …, *P*_*N*_ denote the training subjects.

## 4. Discussion

In this paper, we have explored the use of kernel methods for the prediction of individual traits from EEG spectrograms; and we have introduced KMER, a prediction method based on the idea of interpreting channel spectrograms as probability distributions. By doing this, we could leverage mathematical principles from kernel learning. Although not pursued in this paper, the same principles can also be applied to other problems, such as unsupervised clustering of subjects from EEG spectral information. Even though the mathematical foundations of the method are not particularly trivial, KMER is simple to implement and use, provides spatial interpretation at the sensor or source space, and it is not computationally costly in comparison to more complex (e.g. deep learning) approaches that are estimated on the raw data. The fact that it only necessitates the spectrograms has additional practical benefits in terms of data sharing. These benefits are shared, nonetheless, by KRR.

We tested the methods in the prediction of age in the HarMNqEEG dataset, a recently published international initiative that made EEG spectrograms available across different cohorts and countries. In these data, we observed that KMER and KRR performed similarly, although they both outperformed non-kernelised regression (RR), highlighting the benefits of kernel methods in this context. We note that, compared to other works that predicted age within a single cohort, predicting across sites (i.e. such that models were trained on some sites and tested in others) presents a more challenging problem because the distribution of age across sites varies substantially and potentially also because of protocol or infrastructure differences between the sites. Despite these difficulties, the presented results are comparable to previous EEG studies; see Subsection 2.5. Furthermore, when training and testing within one site only (New York, with a large number of subjects and the broadest age range), the accuracy for some of the individual sensors attained the state-of-the-art accuracies reported in the literature, some of which used more complex models and whole-brain data.

An overarching limitation of all the methods is that, while differences at younger ages are well-predicted, their predictive capacity is much reduced for the older age ranges. A conceivable possibility is that the ageing process is comprised of two separate components of change: a developmental and an ageing part, with more salient differences within development. To investigate this, we performed predictions on the logarithm of the age, which improved results moderately, and we carried out a post hoc bias correction (Smith et al. (2019)), which did improve the results substantially. While the cause of this phenomenon may be important, it falls out of the scope of this work and will require further investigation.

Although we did not exhaustively explored it here, KMER could be further optimised by expanding the selection of the kernel functions and hyperparameters. For instance, while for KMER we optimised for the kernel hyperparameter that generates the kernel matrix from the distance matrix (*f* (·, ·), Equation 4), the kernel used in the construction of the MMD metric (*k*(·, ·), Equation 3) also has a kernel hyperparameter which we set to 1.0 by default. Improving the hyperparameter tuning thus may be a potential avenue for KMER to outperform the simpler KRR approach. Future work will also explore extensions for multichannel predictions and prediction in source spaces.

Finally, it is worth noting that, although we here demonstrated the performance of the tested methods on EEG, KMER (as well as KRR) can be applied to other modalities such as MEG or ECoG; and can be used to predict other individual traits besides age, including cognitive and clinical variables.

## Acknowledgments

D. Vidaurre is supported by a Novo Nordisk Foundation Emerging Investigator Fellowship (NNF19OC-0054895), an ERC Starting Grant (ERC-StG-2019-850404), and a DFF Project 1 from the Independent Research Fund of Denmark (2034-00054B). This research was funded in part by the Wellcome Trust (215573/Z/19/Z). For the purpose of Open Access, the author has applied a CC BY public copyright licence to any Author Accepted Manuscript version arising from this submission. We acknowledge support from PICT 2020-01413. We also thank Sonsoles Alonso for her help.

## Data Availability Statement

The data that support the findings of this study are available in https://www.synapse.org/ with id: *syn*26712693. Complete public access is available by registering and logging into the system.

