## Supplementary material for "Predicting subject traits from brain spectral signatures: an application to brain ageing": SI

| **Country** | **Dataset sites** | **Synapse Name** | **N individuals (Female; Male)** | **Age range** | **Device** |
|---|---|---|---|---|---|
| Barbados | Barbados_1978 ([1]) | Barbados | 62 (F28; M34) | 5.5–11.4 | DEDAAS |
| China | Chengdu_2014 ([2]) | Chengdu | 33 (F7; M26) | 21–28 | BrainAmp |
| | Chongqing_2016 ([3]) | Chongqing | 235 (F134; M101) | 15–26 | BrainAmp |
| Colombia | Colombia_2019 | Colombia | 21 (F13; M8) | 22–45 | Neuro scan |
| Cuba | Cuba_90 ([4]) | Cuba90 | 195 (F98; M97) | 5.5–97 | Medicid-3M |
| | Cuba_2003 ([5]) | Cuba2003 | 48 (F28; M20) | 5–69 | Medicid-4 |
| | Cuba_2004 (?) | Cuba2004 | 14 - | 22–48 | ? |
| | CHBMP ([6]) | CHBMP | 124 (F27; M97) | 17–62 | Medicid-5 |
| Germany | Germany_2013 ([7]) | Germany | 178 (F113; M65) | 22.5–77.5 | BrainAmp |
| Malaysia | Malaysia_2017 | Malaysia | 26 (F24; M2) | 19–60 | ANT Neuro |
| Russia | Russia_2013 ([8]) | Russia | 58 (F34; M24) | 18–49 | nvx136 |
| | | same | 145 (F70; M75) | 16–57 | actiCHamp |
| Switzerland | Bern_1980 ([9]) | Bern | 44 (F18; M26) | 10–16 | Nihon Koh |
| | Zurich_2017 ([10]) | Switzerland | 165 (F80; M85) | 18–90 | EGI-256 HC |
| USA | New York_1970s ([11]) | NewYork | 230 (F109; M121) | 6–80.5 | DEDAAS |
| | Total | 1564 (F783; M781) | | | |

Table 1: Multi-national EEG norms dataset as described in [12]. It consist of 9 countries, 12 devices and 14 batches.

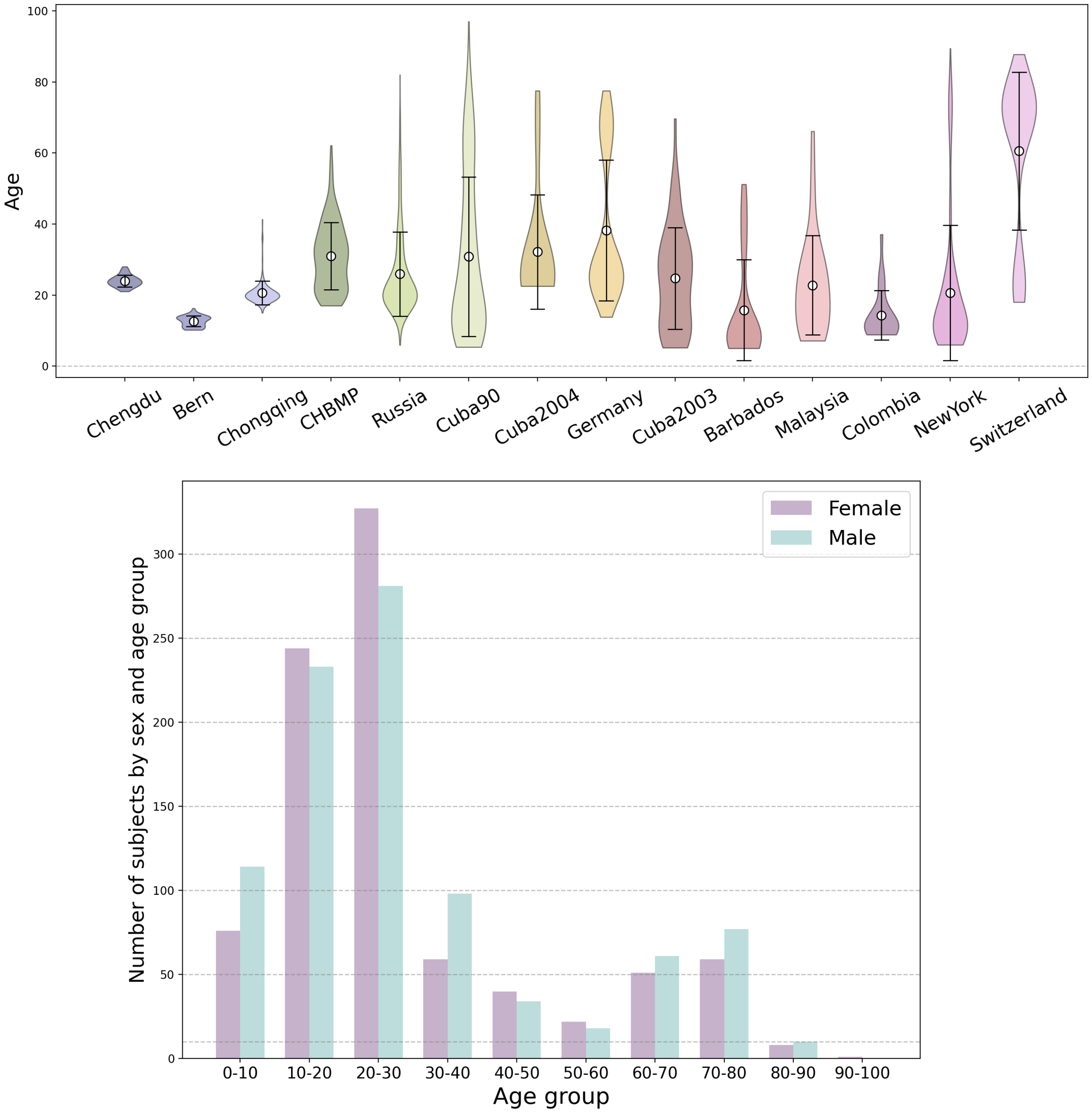


Figure 1: HarMNqEEG dataset demographics. Top Panel: Distribution of individuals in the data set by age and batch. Bottom Panel: Distribution of individuals in the data set by sex and age group. Both figures show that the data set is not uniform in each site or age range.

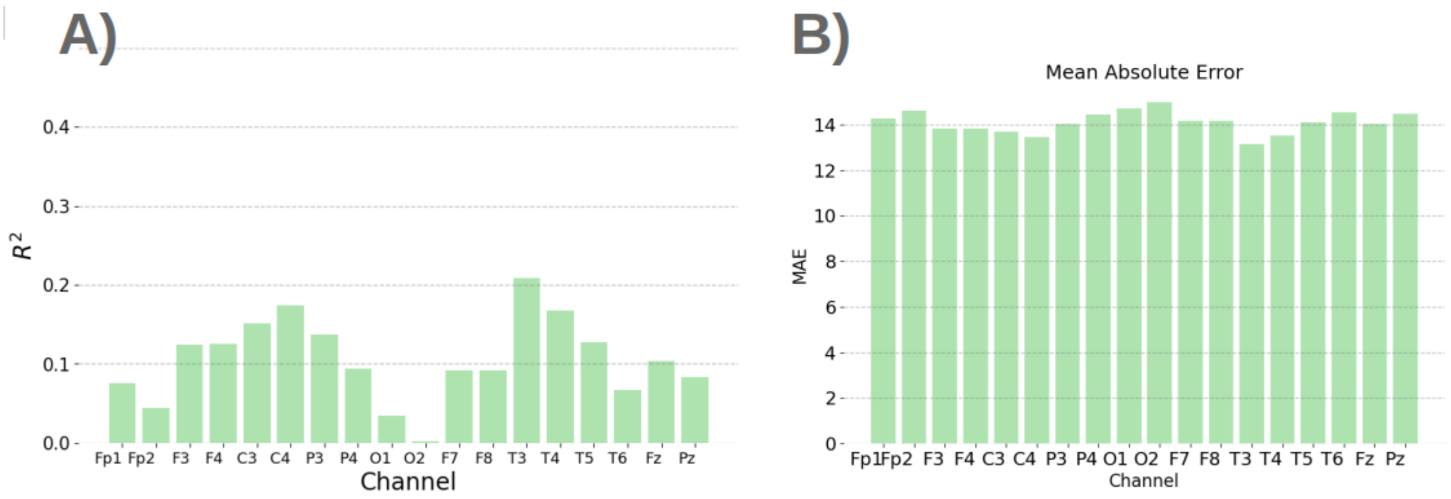


Figure 2: Results for KMER with a linear kernel $f(\cdot)$. **A)** $R^2$. **B)** Mean Absolute Error.

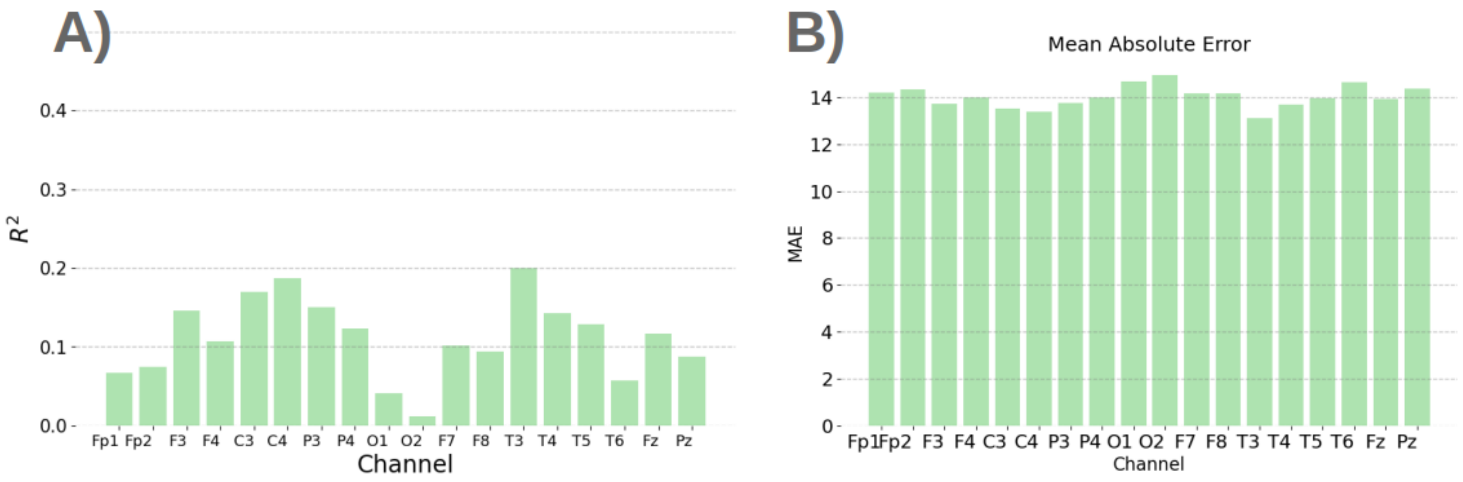


Figure 3: Results for KMER with a polynomial kernel $f(\cdot)$. **A)** $R^2$. **B)** Mean Absolute Error.

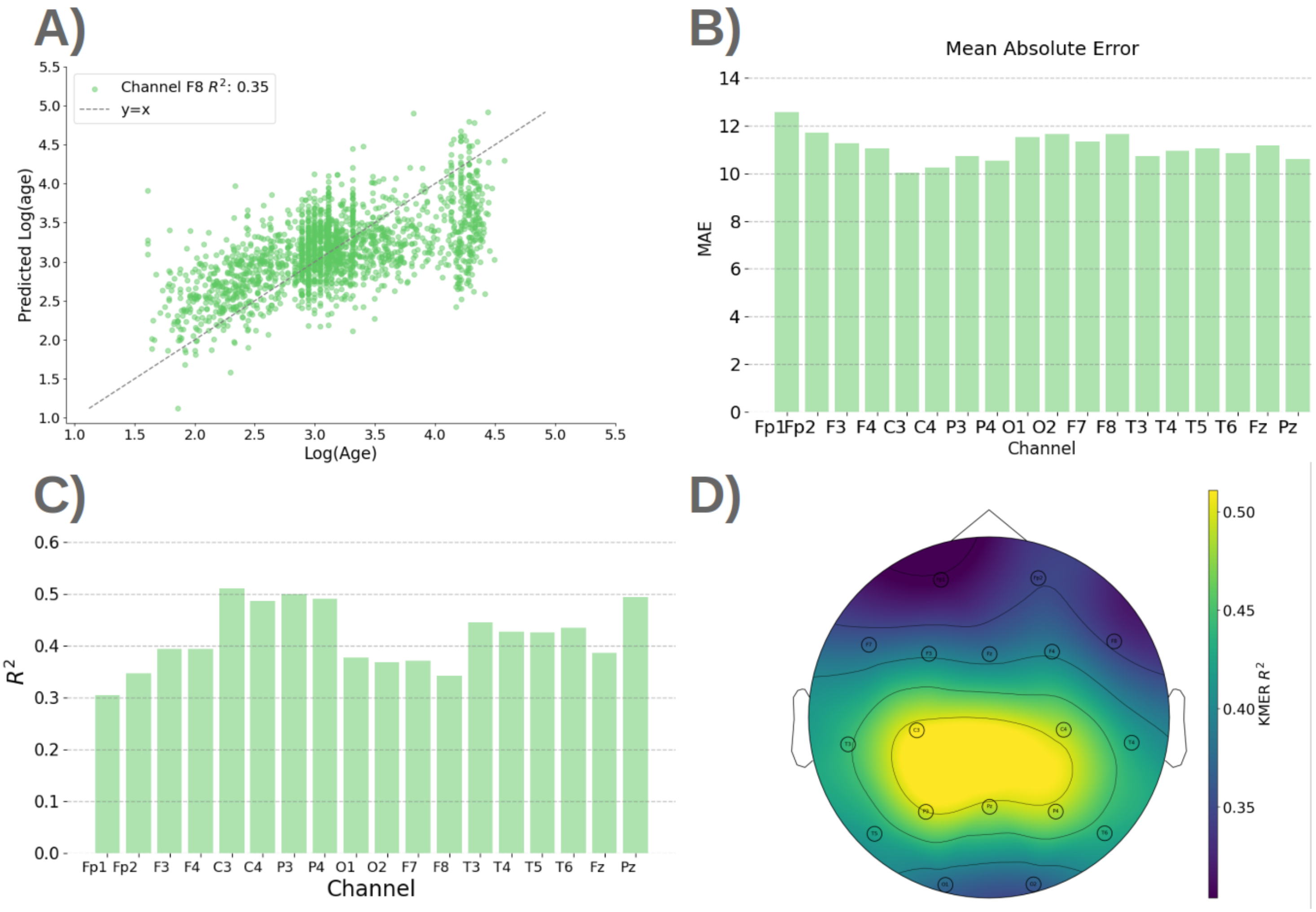


Figure 4: Results for KMER using the logarithmic of age. **A)** Illustration of predicted vs. real age for channel T3. **B)** MAE. **C)** $R^2$. **C)** $R^2$ on a topographic map.

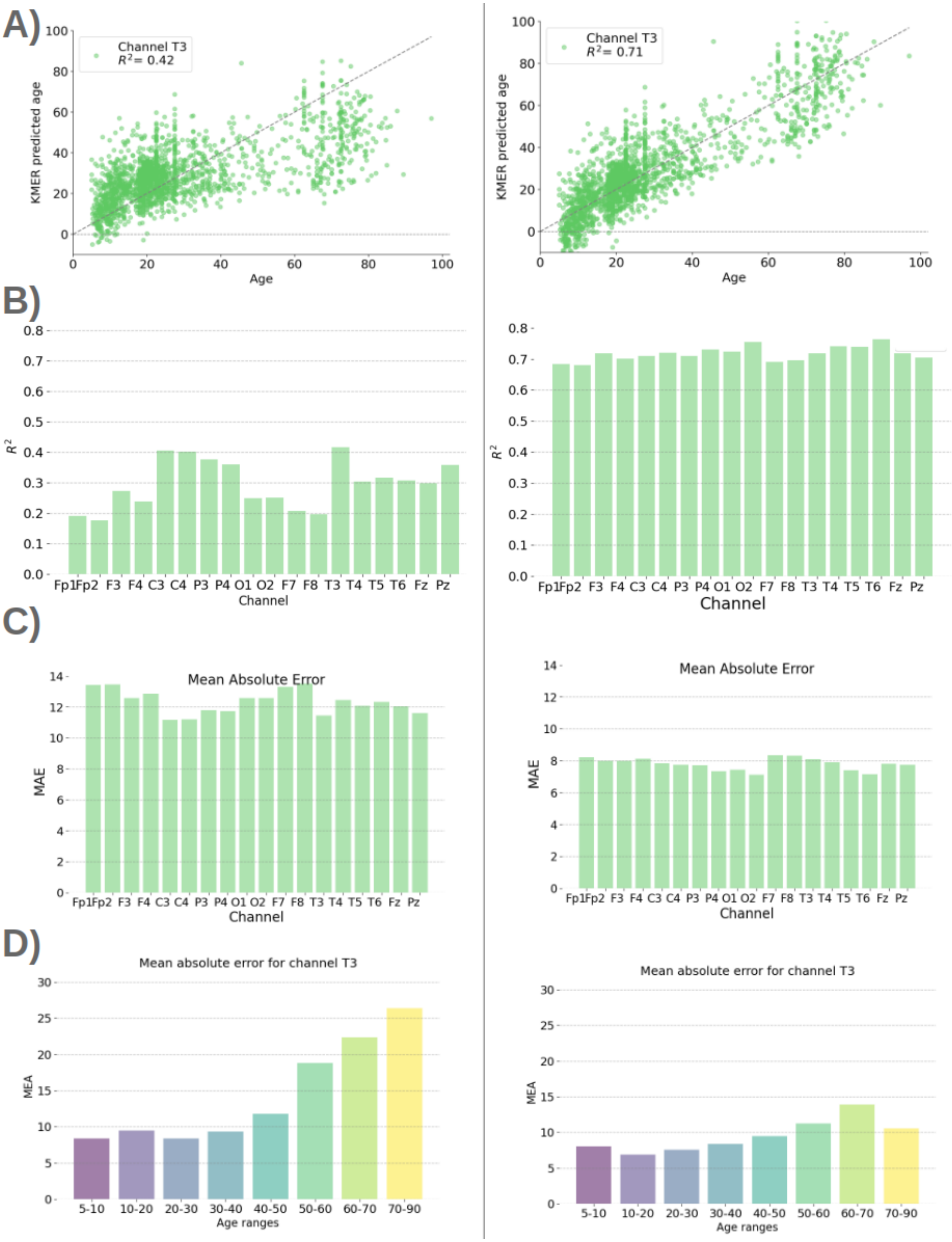


Figure 5: Effect of applying a posthoc bias correction on KMER. Left panels correspond to uncorrected results, and right panels to corrected results. **A)** Illustration of predicted vs. real age for channel T3. **B)** Explained variance $R^2$ per channel. **C)** MAE per channel. **D)** MAE per age range.

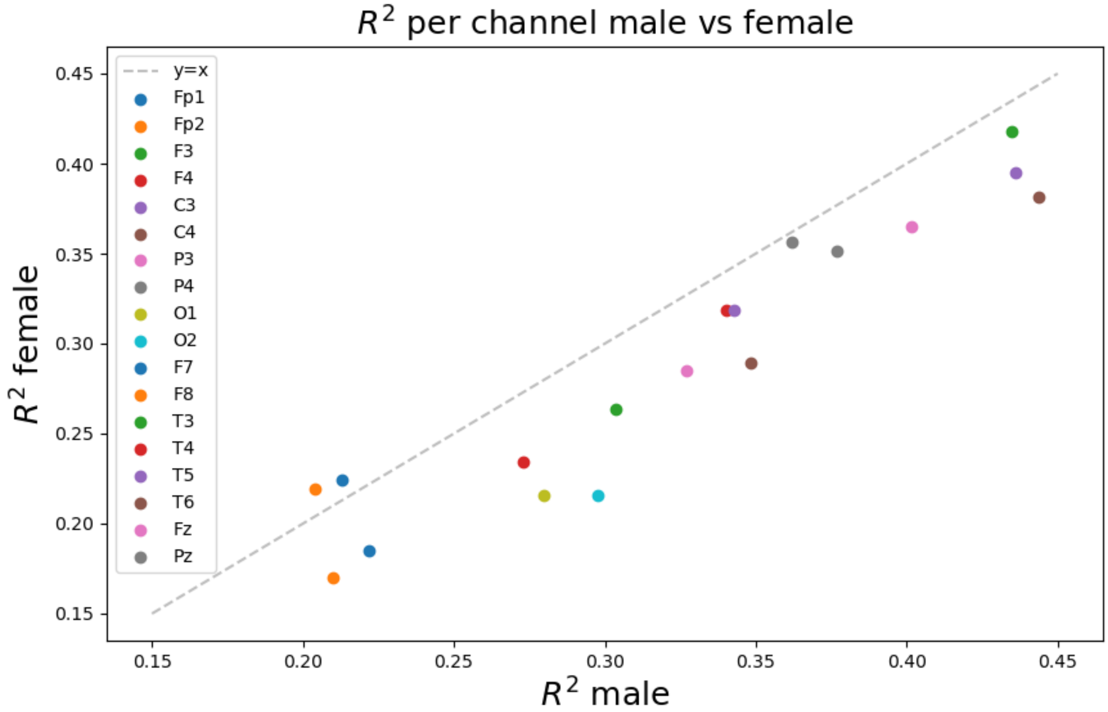


Figure 6: Male vs female accuracies (one point per channel.

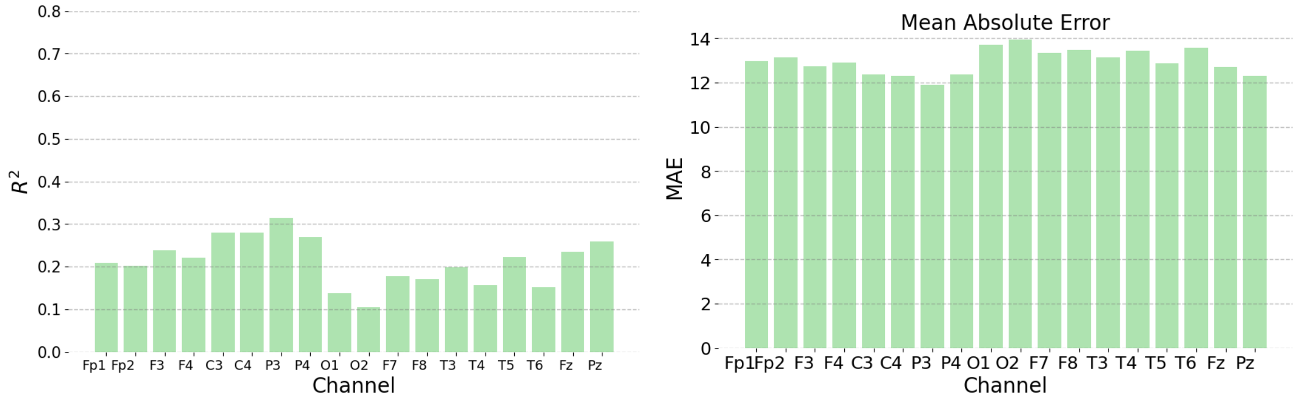


Figure 7: Effect of Normalization. $R^2$ and MAE for KMER with each bin normalize across subjects (before scaling them to sum to 1) such that the variance is comparable across bins.
